# An opposing self-reinforced odor pre-exposure memory produces latent inhibition in *Drosophila*

**DOI:** 10.1101/2021.02.10.430636

**Authors:** Pedro F. Jacob, Paola Vargas-Gutierrez, Zeynep Okray, Stefania Vietti-Michelina, Johannes Felsenberg, Scott Waddell

## Abstract

Prior experience of a stimulus can inhibit subsequent acquisition or expression of a learned association of that stimulus. However, the neuronal manifestations of this learning effect, named latent inhibition (LI), are poorly understood. Here we show that odor pre-exposure produces LI of appetitive olfactory memory performance in *Drosophila*. Behavioral expression of LI requires that the context during memory testing resembles that during the odor pre-exposures. Odor pre-exposure forms an aversive memory that requires dopaminergic neurons that innervate the γ2α′1 and α3 mushroom body compartments - those to α3 exhibit increasing odor-driven activity with successive pre-exposures. In contrast, odor-specific responses of the corresponding mushroom body output neurons are suppressed. Odor pre-exposure therefore recruits specific dopaminergic neurons that provide teaching signals that attach negative valence to the odor itself. LI of *Drosophila* appetitive memory consequently results from a temporary and context-dependent retrieval deficit imposed by competition with this short-lived aversive memory.

## Introduction

Keeping track of life experience allows animals to benefit from all of their prior knowledge when learning new information and using their memory to direct behavior. Although the subject of great early debate among learning theorists, it is now accepted that learning occurs even without explicit rewards or punishment. Classic experiments showed that rats given the prior opportunity to roam in an empty maze performed better when they were later trained with rewards presented in specific locations ^1^. Becoming familiar with the maze without explicit reinforcement and an obvious initial change in the animal’s behavior was called ‘latent learning’ ^2^.

Attempts to replicate a facilitating effect of latent learning using classical conditioning led to an unexpected observation. Pre-exposing animals to a stimulus instead often inhibited the ability of the animal to learn using that stimulus – a phenomenon given the name ‘latent inhibition’ (LI) ^3,4^. LI has been heavily studied for the last 50 years and two alternative theories have been proposed to account for the inhibitory effect of stimulus pre-exposure. In the A, or Acquisition model, subsequent learning is considered to be impaired because pre-exposure alters the capacity for the stimulus to enter into new associations ^5–7^. In contrast in the R, or retrieval, model, learning is still believed to occur but memory expression is impaired ^8–11^. A strong argument in favor of the R model is the observation that LI often appears to be limited in time, leading to expression of the subsequent learning undergoing ‘spontaneous recovery’. Importantly, both theories of latent inhibition assume that something is learned during pre-exposure, such as primitive properties of the stimulus, including its specific identity, intensity (eg. concentration), and salience. In addition, LI is often sensitive to the consistency of the context within which the animal is pre-exposed, taught and tested for memory expression. This led to the proposal that first learning an association between the stimulus and its context makes it difficult for the animal to subsequently associate the stimulus with reinforcement during training ^12,13^.

Studying olfactory learning in the relatively small brain of *Drosophila* has potential to define how LI can operate and reveal the underlying neuronal circuit mechanisms. Several earlier studies in both adult flies and larvae demonstrated that repeated exposure to an odor can alter its apparent valence to the fly, either making the fly avoid it more, or become unresponsive to it ^14–17^. Although LI has been reported with appetitive conditioning in the honeybee ^18,19^, a prior study in adult *Drosophila* did not observe any effect on aversive conditioning following a single odor-pre-exposure ^20^.

Associative olfactory learning in *Drosophila* relies on the neuronal circuitry of the mushroom body. Individual odors are represented as activity in sparse and largely non-overlapping subpopulations of the ~4,000 intrinsic neurons called Kenyon cells (KCs). Positive or negative valence can be assigned to these odor representations by anatomically discrete dopaminergic neurons (DANs) which, via dopamine receptor directed cAMP-dependent plasticity ^21–27^, modulate the efficacy of KC output synapses onto different downstream mushroom body output neurons (MBONs), whose dendrites occupy the same MB compartment. Aversive learning depresses KC synapses onto MBONs whose activation favors approach, whereas appetitive learning reduces odor-drive to MBONs favoring avoidance. By establishing a skew in the valence of the odor-driven MBON network, learned information subsequently directs either odor avoidance or attraction behavior ^28^.

A number of studies indicate that discrete experience is represented as plasticity of different combinations of KC-MBON connections, directed by the engagement of unique combinations of DANs. For example, different types of DANs have been implicated in coding memories for specific rewards, e.g. water, the sweet taste and and nutrient value of sugars, the absence of expected shock and the delayed recognition of safety ^29–34^. In contrast, the same PPL1 DANs appear to be required to code aversive memories for electric shock, bitter taste and heat ^35–37^, although imaging suggests they are activated by a decrease, and not an increase, of temperature ^36,38^. By forming and storing conflicting and complementary memories in different places, the fly can more effectively direct its behavior to reflect a history of prior experience.

Here we show that prior odor exposure temporarily inhibits performance after subsequent appetitive learning in *Drosophila*. This inhibitory effect is sensitive to a change of context across the pre-exposure, training and testing periods, consistent with it being a form of LI. Odor-preexposure forms a short-lived odor-specific aversive memory, whose acquisition requires the γ2α′1 and α3 DANs, the latter of which become sensitized to consecutive odor presentation. As a consequence, aversive memory is apparent as a decrease in the odor-evoked response of the corresponding γ2α′1 and α3 MBONs. Blocking the α3 MBONs impairs the expression of the aversive odor pre-exposure memory and abolishes LI of appetitive memory. The short-lived presence of a parallel and uniquely located odor-specific aversive memory therefore temporarily inhibits the retrieval of a subsequently formed appetitive memory for that same odor. These data provide evidence for a context-dependent R model of latent inhibition in *Drosophila*.

## Results

### Odor pre-exposure can produce temporary latent inhibition of appetitive memory

*Drosophila* can be appetitively conditioned by pairing odor presentation with sugar reward ^15,39^. We therefore tested whether prior exposure to the to be sugar-paired odor (CS+) altered subsequent learned behaviour. Starved flies were given two 2 min odor X presentations with a 15 min intertrial interval (ITI) before being trained by presenting odor Y for 2 min then 30 s later presenting odor X with sucrose reward. Control flies were twice pre-exposed to the odor diluent of mineral oil (MO) before training. When immediately tested after training mineral oil exposed flies exhibited appetitive memory for odor X. However, the performance of flies pre-exposed to the conditioned odor X was significantly impaired (Fig. 1A), a phenotype resembling LI. Since LI can typically be reduced by changing the context between pre-exposure, training and testing, we next altered the context between these experimental stages. In our intitial experiments (Fig. 1A), the flies were pre-exposed in clear tubes, trained in tubes lined with filter paper, and then tested in clear tubes. We next pre-exposed and trained the flies in tubes lined with filter paper to make the context consistent during these stages. This constancy of pre-exposure and training context appeared to slightly enhance the inhibitory effect of pre-exposure (Fig. 1B). More importantly, these first two experiments suggest that flies may consider the context of clear (A) and paper (A*) tubes to be similar. We next explicitly changed the context by pre-exposing flies to the to be CS+ in a copper grid-lined tube (B) before training them in tubes lined with filter paper (A*), and testing them in clear tubes (A) (Fig. 1C). Strikingly, the robust change in context between copper grid tubes and paper/clear (BA*A) abolished the inhibitory effect of odor pre-exposure. A failure to observe LI across changing context could result from the inability of the flies to form, or retrieve a pre-exposure memory. However, robust LI was recovered when the flies were trained on paper but were pre-exposed and tested in copper grid tubes (BA*B) (Fig. 1D), showing that pre-exposure memory is formed but not retrieved in the BA*A scenario. Together these context-shifting experiments demonstrate that the inhibitory effect of odor pre-exposure on subsequent appetitive memory is a form of LI, whose expression depends on the context at testing approximating that experienced during odor pre-exposure.

**Figure 1.**
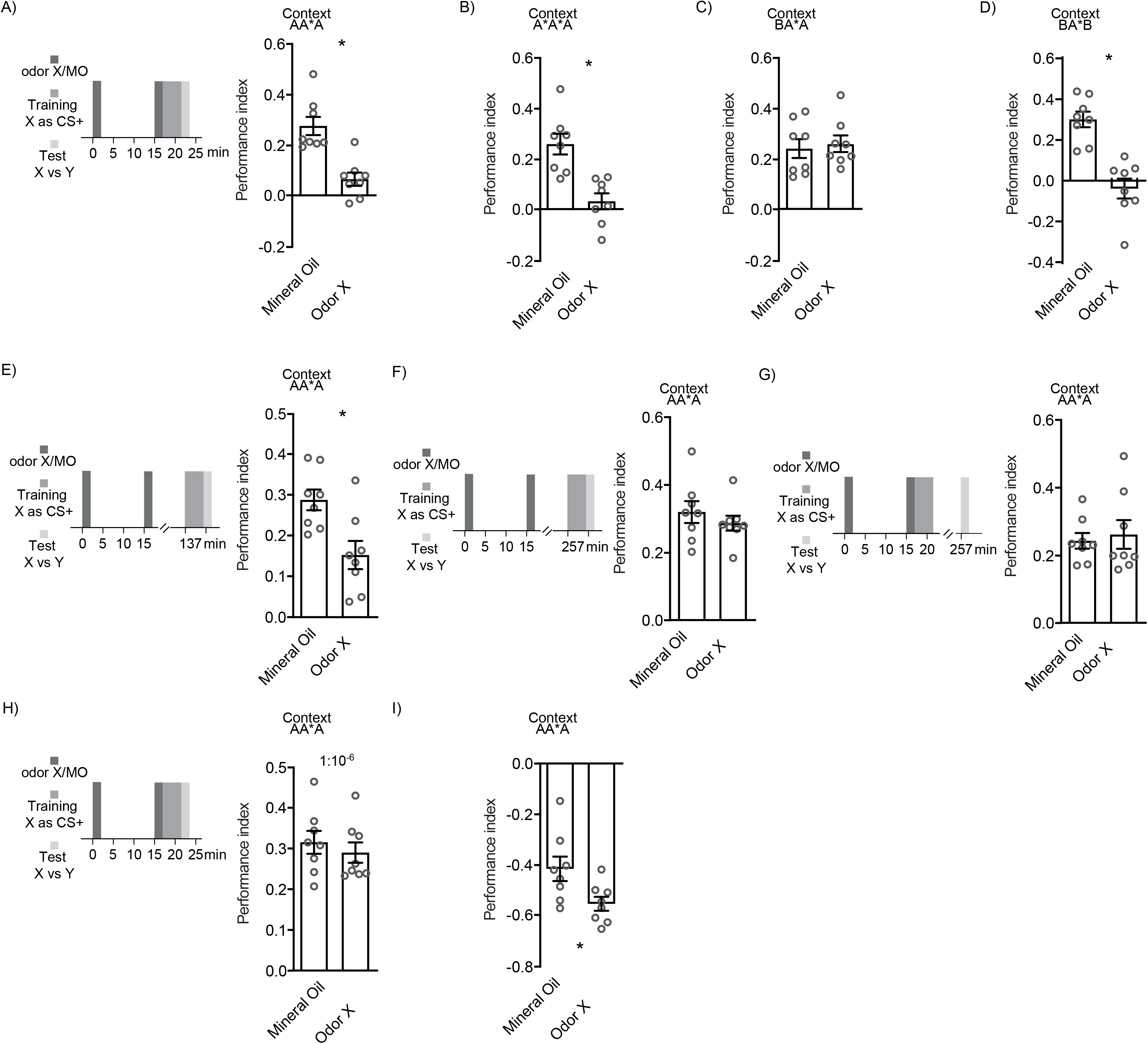
Odor pre-exposure induces context-dependent latent inhibition of appetitive learning. (**A**) Flies were twice pre-exposed to odor X or mineral oil (MO) in clear tubes (context A), with a 15 min inter-trial interval. Immediately following the last exposure, they were trained by presenting odor Y for 2 min, then odor X for 2 min with sugar (CS+; context A*). Flies pre-exposed to odor X exhibited appetitive memory performance to odor X (one-sample t-test: t(7)=2.419, p=0.0461) that was significantly reduced (LI) in comparison to flies exposed to MO [t(14)=4.719, p=0.0003]. (**B**) When pre-exposure and training context were more closely matched, using filter paper in the pre-exposure tube (context A*A*A), flies pre-exposed to odor X did not exhibit appetitive memory performance (LI) (one-sample t-test: t(7)=0.9866, p=0.3567). Performance was also significantly reduced compared to flies pre-exposed to MO [t(14)=4.376, p=0.0006]. (**C**) Pre-exposing flies to odor X in a different context of a copper grid tube (context BA*A) abolished the inhibitory effect (no LI). Appetitive memory performance of flies pre-exposed to odor X was similar to MO pre-exposed flies [t(14)=0.3902, p=0.7023]. (**D**) Matching context between pre-exposure and testing (both in copper grid tube; context BA*B) restored LI. Performance of flies pre-exposed to odor X was significantly impaired compared to flies pre-exposed to MO [t(14)=5.449, p<0.0001]. (**E**) The inhibitory effect on appetitive learning is evident 2 h after odor pre-exposure. Flies were pre-exposed to Odor X or MO then 2 h later were appetitively trained and tested immediately for memory. Performance of flies pre-exposed to odor X was significantly reduced compared to flies pre-exposed to MO [t(14)=3.155, p=0.007]. (**F**) The inhibitory effect on learning was not evident 4 h after odor pre-exposure. Memory performance of flies pre-exposed to odor X or MO was similar when trained 4 h after the last pre exposure [t(14)=0.8368, p=0.4167]. (**G**) The inhibitory effect of pre-exposure on learning was not evident 4 h after training. Performance was similar if flies were pre-exposed to odor X or MO, then immediately trained but tested for memory 4 h later [t(14)=0.3877, p=0.7041], demonstrating time-dependent loss of LI and recovery of appetitive memory. (**H**) Pre-exposure to lower odor concentration did not induce LI. Flies were pre-exposed with 1:10^−6^ odor X or mineral oil (MO), with 15 min inter-trial interval, in clear tubes, immediately trained with 1:10^−6^ odor X paired with sugar (CS+) and tested for memory. Flies pre-exposed to 1:10^−6^ odor X showed similar memory performance to those pre-exposed to MO [t(14)=0.6741, p=0.5112]. (**I**) Odor pre-exposure facilitates memory performance after aversive conditioning. Flies were pre-exposed to odor X or mineral oil (MO) in a copper grid-lined tube (context BBA), then immediately trained by pairing odor X for 1 min with twelve 30V electric shocks (CS+). Flies pre-exposed to odor X exhibited increased aversive memory performance in comparison to MO-exposed flies [t(14)=2.485, p=0.0262].

We next tested the persistence of LI. Whereas pre-exposure to the CS+ impaired memory performance when flies were trained 2 h after the last pre-exposure (Fig. 1E), performance was unimpaired if training was delayed for 4 h after the last pre-exposure (Fig. 1F). In addition appetitive memory performance was unimpaired if flies were trained immediately after the second pre-exposure but tested 4 h later (Fig. 1G). These experiments therefore suggest that the inhibitory effect of LI decays between 2 and 4 h after odor pre-exposure. In addition, the recovery of appetitive memory performance is consistent with LI resulting from a temporary retrieval deficit.

Many studies have shown that LI is sensitive to properties (frequency, amount and duration) of the conditioned stimulus ^40–42^. We therefore also tested whether two exposures of a lower odor concentration (10^−6^ rather than 10^−3^) with 15 min ITI induced LI of appetitive memory. Although training with this odor concentration produced robust appetitive memory, no LI effect was observed following odor pre-exposure (Fig. 1H).

### Odor pre-exposure can produce facilitation of aversive memory

We also tested whether odor pre-exposure altered performance measured after aversive training, pairing odor with electric shock. Starved flies were given two 2 min odor X presentations with a 15 min ITI before being trained by presenting odor X for 1 min, paired with electric shocks, then 45 s later presenting odor Y. Surprisingly, flies pre-exposed to the CS+ exhibited faciliated aversive memory performance as compared to flies pre-exposed to mineral oil (Fig. 1I). Finding that the same odor pre-exposure schedule can inhibit appetitive memory but faciliate aversive memory performance led us to hypothesize that pre-exposure might form an avoidance memory for the CS+, that in the former case is competing and the latter complementary.

### Pre-exposure forms a short-lived mushroom body-dependent aversive odor memory

Prior work has shown that odor pre-exposure can enhance subsequent odor avoidance behavior ^14,15,33^, consistent with our hypothesized aversive learning model. We therefore tested whether such an effect resulted following our LI odor exposure regimen. As in the prior experiments starved flies were exposed twice to 2 min of odor X (10^−3^ dilution in mineral oil) with a 15 min ITI. They were then immediately tested for preference between the pre-exposed odor X and another odor Y, without any training. Flies pre-exposed twice to odor X showed a selective avoidance of odor X, consistent with pre-exposure forming an aversive odor memory (Fig. 2A). In contrast, a single odor pre-exposure, of either 2 or 4 min, did not alter odor preference, suggesting that repeated exposure is required to form the avoidance memory (Fig. 2A). Measuring odor preference at different times after pre-exposure revealed that the avoidance memory is labile and slowly decays between 15 min and 2 h (Fig. 2B). We also tested whether expression of pre-exposure memory was sensitive to context by pre-exposing flies to odor in copper grid tubes and testing them for odor preference in clear tubes. Performance was unaffected by this change of context (Fig. 2C), demonstrating that context is uniquely important for the LI effect of pre-exposure memory. We last tested whether the explicit absence of food acts as aversive reinforcement for hungry flies by pre-exposing satiated flies. However, pre-exposure induced similar odor avoidance in satiated flies as compared to starved flies (Fig. 2D). Since our LI effect was assayed in hungry flies, all subsequent experiments in this study were performed in hungry flies.

**Figure 2.**
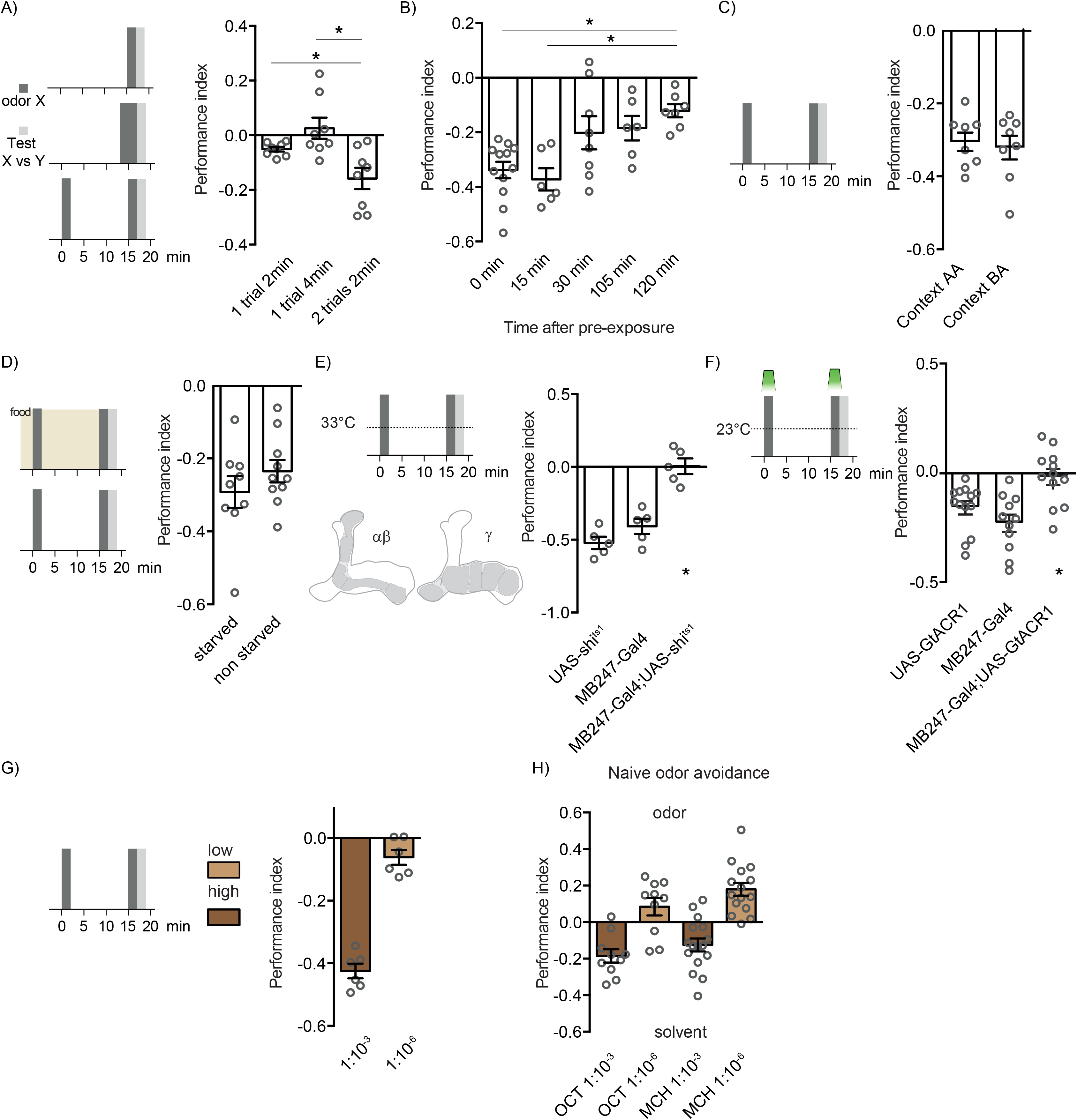
Odor pre-exposure forms a labile mushroom body-dependent aversive memory. (**A**) Repeated odor-exposure forms aversive memory for that odor. Two 2 min odor exposures with 15 min interval induced odor aversion [F(2,21)=8.365, p=0.0023, n=8] whereas a single 2 min or 4 min exposure did not [Tukey’s multiple comparisons test: 1×2 min *vs.* 1×4 min, p=0.07; 1×2 min *vs.* 2×2 min, p=0.0182; 1×4 min *vs.* 2×2 min, p=0.0048]. (**B**) Pre-exposure induced aversive memory is labile [one sample t-test p<0.05 for all comparisons; F(4,34)=6.295, p=0.007, n=6-12; Tukey’s multiple comparisons test: 0 min *vs.* 120 min, p=0.0036 and 15 min *vs.* 120 min, p=0.0040, for all other comparisons p>0.05). (**C**) Expression of pre-exposure memory is not sensitive to changing context. Performance of flies pre-exposed in copper-lined and tested in clear tubes (context BA) was indistinguishable from that of flies pre-exposed and tested in clear tubes (context AA) [t(14)=0.3827, p=0.7077]. (**D**) Formation of odor pre-exposure memory does not depend on hunger [t(17)=1.090, p=0.2911]. (**E**) Blocking synaptic output from the MB KCs abolishes odor pre-exposure memory [F(2,12)=30.87, p<0.0001, n=5; Tukey’s multiple comparisons test: UAS-*Shi*^ts1^ *vs.* MB247-GAL4, p=0.2773; UAS-*Shi*^ts1^ *vs.* MB247-GAL4;UAS-*Shi*^ts1^, p<0.0001; MB247-GAL4 *vs.* MB247-GAL4;UAS-*Shi*^ts1^, p=0.0002]. (**F**) Forming pre-exposure memory requires KC activity during odor pre-exposure. Restricting KC block to the period of odor pre-exposure, using GtACR1 and green light, abolishes aversion to odor X [F(2,33)=9.088, p=0.0007, n=11-13; Tukey’s multiple comparisons test: UAS-GtACR1 *vs.* MB247-GAL4, p=0.3423; UAS-GtACR1 *vs.* MB247-GAL4;UAS-GtACR1, p=0.0183; MB247-GAL4 *vs.* MB247-GAL4;UAS-GtACR1, p=0.0006]. (**G**) Formation of pre-exposure memory depends on odor concentration. pre-exposure to high (10^−3^), but not low (10^−6^) concentration of odor induces aversive odor memory [t(10)=10.98, p<0.0001]. (**H**) Reducing odor concentration by 3 orders of magnitude can switch naïve valence from avoidance to attraction. All pre-exposure experiments were performed in the context A (clear tubes) for pre-exposure and testing phases.

Olfactory memories typically depend on the neuronal circuitry of the MB ^39,43–47^. We therefore tested the consequence of blocking output from αβ and γ subsets of MB KCs, on the effect of odor pre-exposure. We expressed the dominant temperature-sensitive UAS-*Shibire*^ts1^ (UAS-*Shi*^ts1^) transgene ^48^ with MB247-GAL4 and blocked KC output throughout the experiment by raising the temperature to restrictive 33°C. This manipulation abolished the development of odor avoidance performance (Fig. 2E). Moreover, restricting inhibition of KC activity to only the periods of odor pre-exposure, using expression of the green light-sensitive anion-selective GtACR1 channel ^49^ revealed that KC activity is necessary during pre-exposure (Fig. 2F). Finding a role for the MB suggests the pre-exposure effect is a form of associative learning.

Since we observed LI when flies were twice pre-exposed to 10^−3^ odor concentration but not to 10^−6^, we also tested whether this lower odor concentration produced an aversive pre-exposure memory. Consistent with the lack of LI, no enhanced avoidance was observed following 10^−6^ odor exposures (Fig. 2G). In addition, when naïve flies were given the choice between a 10^−3^ odor stream and air they exhibited avoidance of the odor (Fig. 2H). In contrast, they either showed no preference, or odor approach when tested with a 10^−6^ odor stream and air. We therefore reasoned that lower odor concentrations, which are less repellent ^50^, do not act as an aversive reinforcer during odor pre-exposure.

### Pre-exposure memory requires γ2α′1 and α3 DANs

Aversive olfactory learning reinforced by electric shock, bitter taste or heat depends on punishment coding DANs from the PPL1 cluster ^35,37,51–54^. In contrast, DANs in the PAM cluster mostly code for reward ^29,30,34^. We therefore first used TH-GAL4 and R58E02-GAL4 to express UAS-*Shi*^ts1^ and test the respective roles of PPL1 and PAM DANs in pre-exposure learning. Blocking TH-GAL4 neurons switched the effect of pre-exposure from generating aversion to approach (Fig. 3A). This reversal of valence implies that removing aversive signalling may either release (mutually exclusive) or unmask (in parallel) positive reinforcement induced by odor pre-exposure. Blocking the rewarding DANs with R58E02-GAL4; with UAS-*Shi*^ts1^ increased the aversive effect of pre-exposure (Fig. 3B), consistent with odor exposure engaging negative and positive reinforcing DAN populations in parallel.

**Figure 3.**
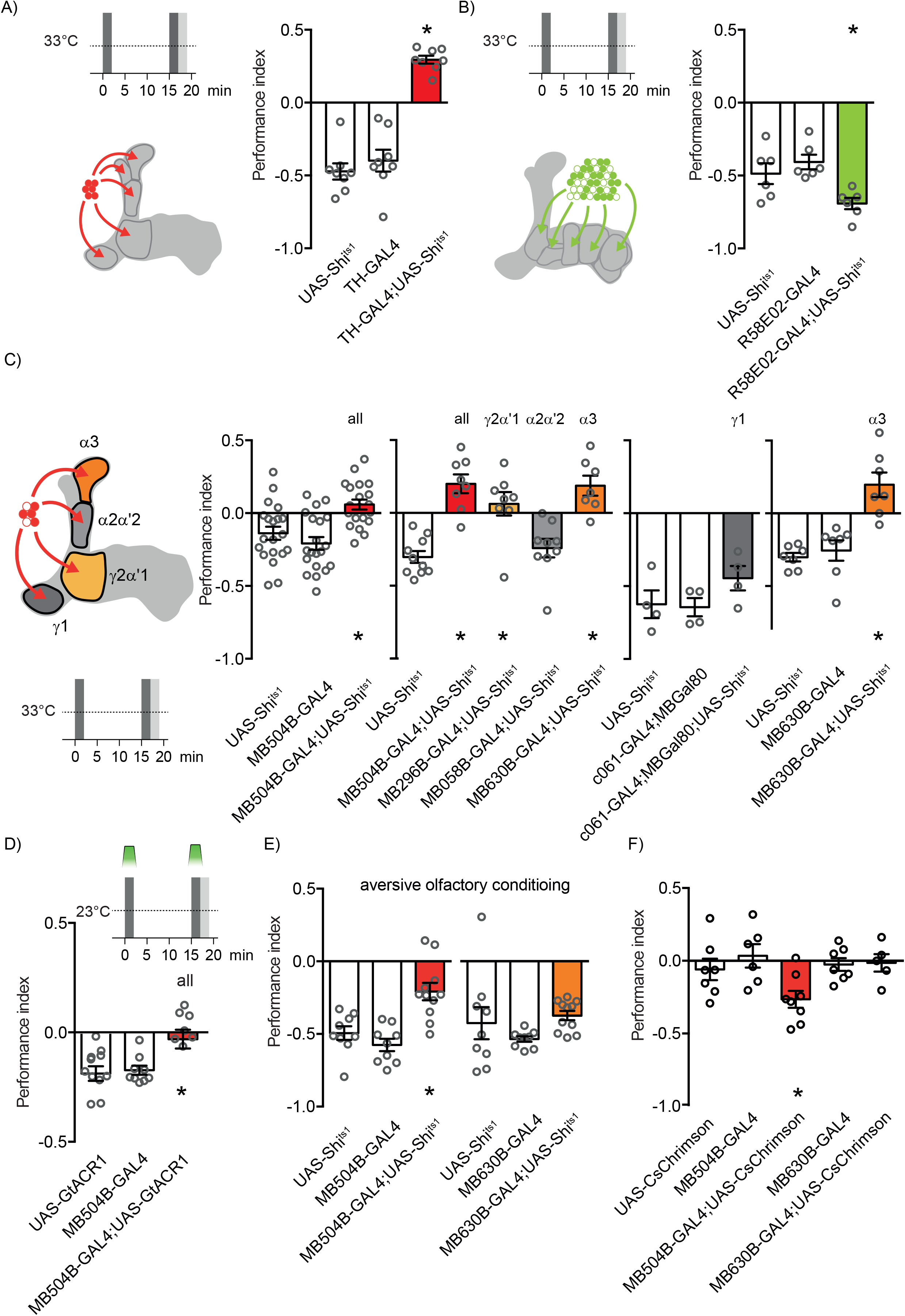
Forming pre-exposure memory requires γ2α′1 and α3 dopaminergic neurons. (**A**) Blocking punishment coding DANs switches the valence of odor pre-exposure learning. Exposure induced aversion becomes approach [F(2,21)=57.26, p<0.0001, n=8; Tukey’s multiple comparisons test: UAS-*Shi*^ts1^ *vs.* TH-GAL4, p=0.6528;UAS-*Shi*^ts1^ *vs.* TH-GAL4;UAS-*Shi*^ts1^, p<0.0001; TH-GAL4 *vs.* TH-GAL4; UAS-*Shi*^ts1^, p<0.0001]. (**B**) Blocking reward coding DANs increases the aversive effect of odor pre-exposure [F(2,15)=7.126, p=0.0067, n=6; Tukey’s multiple comparisons test: UAS-*Shi*^ts1^ *vs.* R58E02-GAL4, p=0.5676; UAS-*Shi*^ts1^ *vs.* R58E02-GAL4/UAS-*Shi*^ts1^, p=0.0473; R58E02-GAL4 *vs.* R58E02-GAL4/UAS-*Shi*^ts1^, p=0.0062]. (**C**) Pre-exposure-evoked aversive learning requires PPL1 DANs [F(2,57)=11.30, p<0.0001, n=20; Tukey’s multiple comparisons test: UAS-*Shi*^ts1^ *vs.* MB504B-GAL4, p=0.4573; UAS-*Shi*^ts1^ *vs.* MB504B-GAL4;UAS-*Shi*^ts1^, p=0.0037; MB504B-GAL4 *vs.* MB504B-GAL4;UAS-*Shi*^ts1^, p<0.0001]. A specific requirement was observed for PPL1-γ2α′1 (MV1) and PPL1-α3 DANs [For PPL1 screen: F(4,37)=14.80, p<0.0001, n=7-10; Dunnett’s multiple comparisons test: UAS-*Shi*^ts1^ *vs.* MB504B-GAL4;UAS-*Shi*^ts1^, p<0.0001; UAS-*Shi*^ts1^ *vs.* MB296B-GAL4;UAS-*Shi*^ts1^, p=0.0006; UAS-*Shi*^ts1^ *vs.* MB058B-GAL4;UAS-*Shi*^ts1^, p=0.8902; ; UAS-*Shi*^ts1^ *vs.* MB630B-GAL4;UAS-*Shi*^ts1^, p<0.0001; For PPL1-γ1: F(2,9)=1.785, p=0.2223, n=4; For PPL1-α3: F(2,29)=23.16, p<0.0001, n=10-11; Tukey’s multiple comparisons test: UAS-*Shi*^ts1^ *vs.* MB630B-GAL4, p=0.5575; UAS-*Shi*^ts1^ *vs.* MB630B-GAL4;UAS-*Shi*^ts1^, p<0.0001; MB630B-GAL4 *vs.* MB630B-GAL4;UAS-*Shi*^ts1^, p<0.0001]. (**D**) Optogenetic silencing of PPL1 DANs during odor exposures blocks exposure-evoked aversive learning [F(2,24)=6.325, p=0.0062, n=9; Tukey’s multiple comparisons test: UAS-GtACR1 *vs.* MB504B-GAL4, p=0.9832; UAS-GtACR1 *vs.* MB504B-GAL4;UAS-GtACR1, p=0.0112; MB504B-GAL4 *vs.* MB504B-GAL4;UAS-GtACR1, p=0.0169]. (**E**) Blocking PPL1 DANs impairs immediate aversive memory following conditioning pairing odor with electric shocks [F(2,26)=14.43, p<0.0001, n=9-11; Tukey’s multiple comparisons test: UAS-*Shib*^ts1^ *vs.* MB504B-GAL4, p=0.5452; UAS-*Shib*^ts1^ *vs.* MB504B-GAL4;UAS-*Shib*^ts1^, p=0.0016; MB504B-GAL4 *vs.* MB504B-GAL4;UAS-*Shib*^ts1^, p<0.0001]. However, only blocking the PPL1-α3 DANs has no effect [F(2,26)=1.692, p=0.2039, n=9-11]. (**F**) Pairing odor exposure with optogenetic activation of PPL1 DANs induces aversive memory to the previously-paired odor, whereas activation of only PPL1-α3 DANs is ineffective [F(4,41)=5.665, p=0.0010, n=7-11; Bonferroni’s multiple comparisons test: UAS-CsChrimson *vs.* MB504B-GAL4, p>0.9999;UAS-CsChrimson *vs.* MB504B-GAL4;UAS-CsChrimson, p=0.0025; MB504B-GAL4 *vs.* MB504B-GAL4; UAS-CsChrimson, p=0.0028; UAS-CsChrimson *vs.* MB630B-GAL4, p>0.9999; UAS-CsChrimson *vs.* MB630B-GAL4;UAS-CsChrimson, p>0.9999; MB630B-GAL4 *vs.* MB504B-GAL4;UAS-CsChrimson, p>0.9999]. All pre-exposure experiments were performed in the context A (clear tubes) for pre-exposure and testing phases.

More specifically blocking the four types of PPL1-DANs implicated in shock-reinforced olfactory learning, either throughout the experiment (Fig. 3C), or only during odor pre-exposure (Fig. 3D) abolished the learned aversion. Blocking the individual types revealed that the PPL1-γ1pedc (MB-MP1) and PPL1-α′2α2 DANs that are important for shock-reinforced olfactory learning ^54^, are dispensable for pre-exposure learning. In contrast, blocking PPL1-γ2α′1 DANs (MB-MV1) abolished pre-exposure learning, while blocking the PPL1-α3 DANs converted pre-exposure-induced aversion to approach (Fig. 3C).

Since PPL1-DANs are thought to provide an aversive teaching signal, common to electric shock, heat and bitter taste we also tested for a role of PPL1-α3 DANs in shock-reinforced aversive memory. Although blocking the four PPL1-DANs impaired shock learning, blocking only the PPL1-α3 DANs left immediate shock memory intact (Fig. 3E). In addition, replacing shock with optogenetic stimulation of the four PPL1-DANs, could artificially implant an aversive memory whereas a single pairing of odor presentation with PPL1-α3 DAN activation did not form aversive memory (Fig. 3F). These data are consistent with a prior study that showed learning requires multiple trials of PPL1-α3 DAN activation ^55^. These results demonstrate that two-trial pre-exposure learning and single-trial electric shock learning involve different PPL1-DANs and emphasize the importance of PPL1-α3 DANs for pre-exposure learning.

### The activity of PPL1-γ2α′1 and PPL1-α3 DANs is altered following odor presentation

Since pre-exposure learning depends on repeated trials and requires the PPL1-γ2α′1 and PPL1-α3 DANs we imaged DAN activity during and after odor presentations under the microscope (Fig. 4). Flies were constructed that expressed the fluorescent UAS-GCaMP6m calcium sensor in PPL1-γ2α′1 or PPL1-α3 or DANs. As before in the behavioral experiments, flies were given two odor presentations with 15 min ITI. They were then immediately given another 5 s exposure of the trained odor followed by 5 s of a novel odor, to mimic the behavioral test situation under the microscope. Odor-evoked responses of PPL1-α3 DANs increased from the first to the second trial (Fig. 4A), whereas PPL1-γ2α′1 DAN responses did not change (Fig. 4C). However, PPL1-γ2α′1 DANs exhibited a reduced response to the trained compared to the novel odor in the 5s test, whereas PPL1-α3 DANs showed no difference between novel and test odor responses (Fig. 4B and 4D).

**Figure 4.**
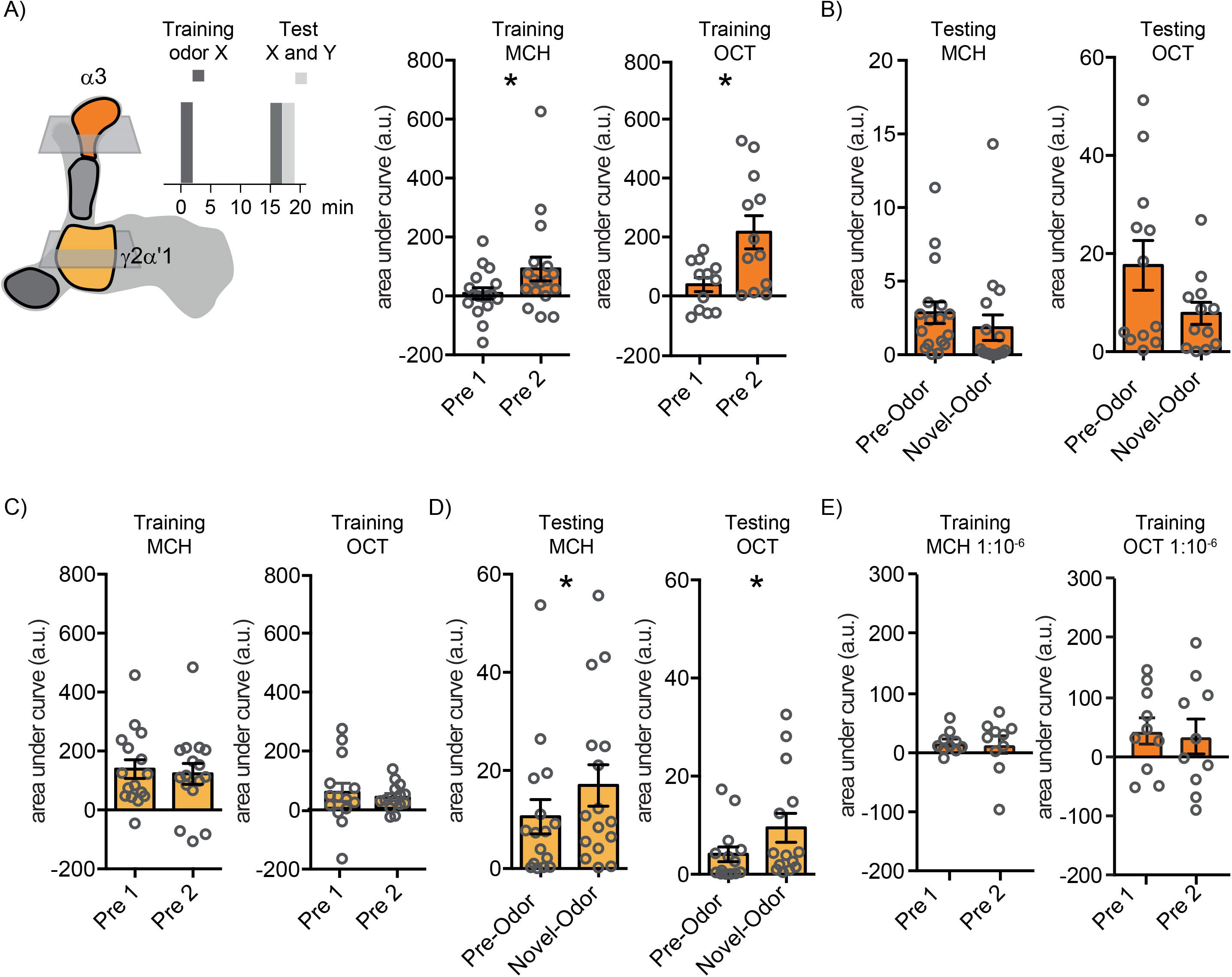
PPL1-γ2α′1 and PPL1-α3 DAN activity is altered following odor exposure. (**A**) Odor-evoked responses of PPL1-α3 DANs increase from the first to the second odor presentation [for MCH: W(18)=91, p=0.0438; for OCT: t(11)=2.859, p=0.0156]. (**B**) In the test trained odor-evoked responses of PPL1-α3 DANs were not different to those for the novel odor [for MCH: W(18)=-33, p=0.4951; for OCT: W(12)=-40, p=0.1294]. (**C**) Odor-evoked responses of PPL1-γ2α′1 DANs did not change between the first and second odor presentations [for MCH: t(17)=0.4792, p=0.6379; for OCT: t(13)=0.5678, p=0.5798]. (**D**) In the test trained odor-evoked responses of PPL1-γ2α′1 DANs were reduced response compared to those for the novel odor [for MCH: W(18)=107, p=0.0182; for OCT: W(14)=71, p=0.0245]. (**E**) Responses of PPL1-α3 DANs did not change when flies were twice exposed to lower concentration of odor [for MCH: t(9)=0.1930, p=0.8513; for OCT: t(9)=0.2901, p=0.7783].

In addition, consistent with a lack of LI (Fig. 1H) and pre-exposure memory (Fig. 2F), lower 10^−6^ odor concentration did not increase the odor-evoked responses of PPL1-α3 DANs (Fig. 4E). Together these results suggest that the increased odor-driven activity of the PPL1-α3 DANs, specifically in the 2^nd^ pre-exposure, is crucial for the formation of an aversive pre-exposure memory.

### Repeated odor presentation depresses odor responses of γ2α′1 and α3 MBONs

The predominant model for Drosophila learning is dopamine-driven depression of synapses between odor-specific KCs and MBONs ^27,56–60^. We therefore used MBON expression of GCaMP6m to test whether KC-MBON connections underlying the PPL1-α3 and PPL1-γ2α′1 DANs were changed following repeated odor exposure (Fig. 5). Odor responses of both MBON-α3 and MBON-γ2α′1 were both reduced compared to their responses to a novel odor (Fig. 5A and 5B). In addition, blocking either the α3 or γ2α′1 MBONs throughout an odor exposure experiment with UAS-*Shi*^ts1^, or specifically during the test phase with UAS-*GtACR1* abolished exposure-induced avoidance behavior in the T-maze (Fig. 5C-5E). Taken together these data indicate that spaced odor exposure forms aversive memory, that manifests as reduced odor-evoked activity of the α3 and MBON-γ2α′1 MBONs resulting from increased odor-evoked activation of the corresponding DANs.

**Figure 5.**
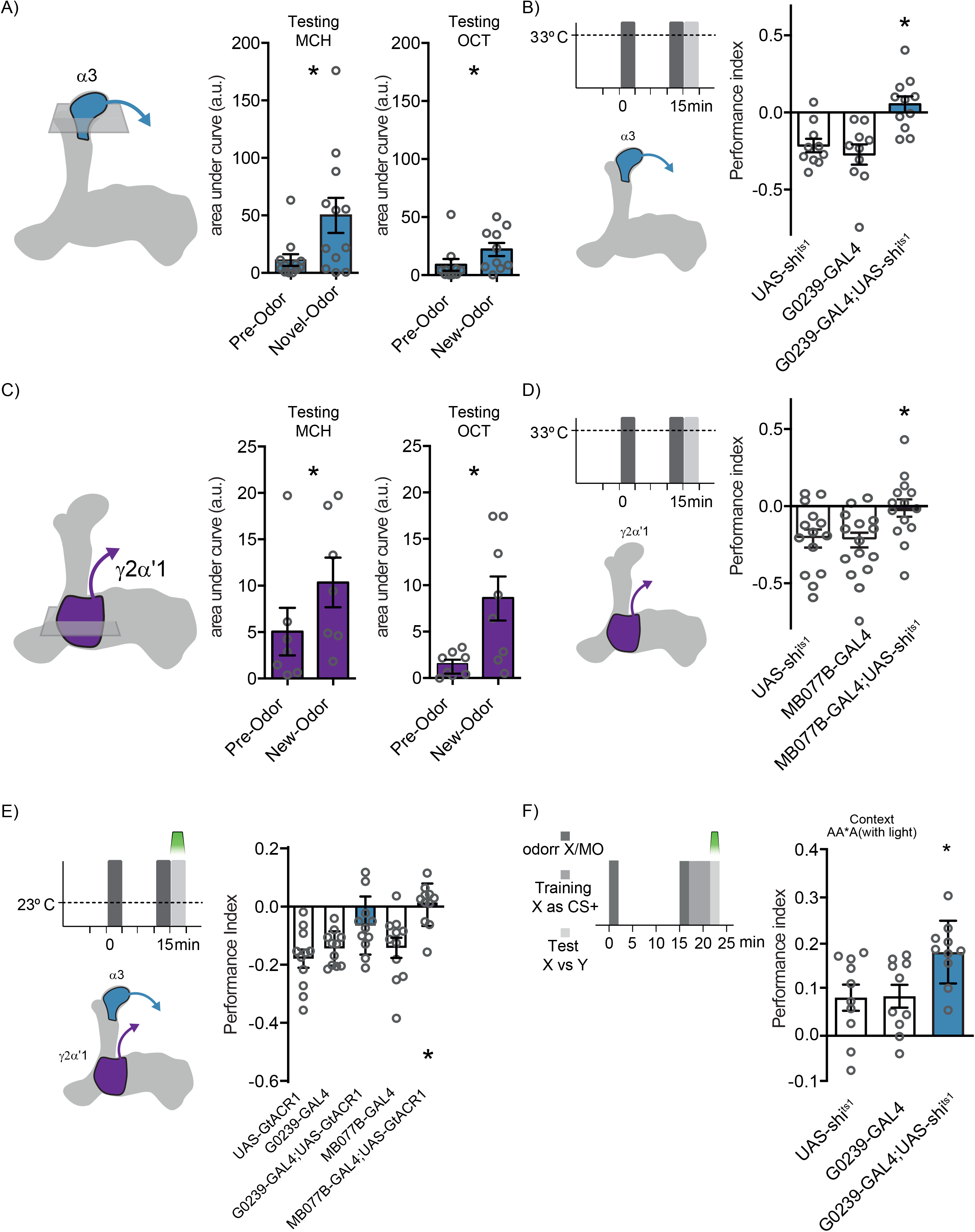
γ2α′1 and α3 MBONs exhibit an odor pre-exposure memory trace and are required for behavioral expression of odor aversion and LI of appetitive memory. (**A**) Odor pre-exposure reduces trained odor responses of MBON-α3 compared to a novel odor [for MCH: W(12)=56, p=0.0269; for OCT: W(13)=57, p=0.0429]. Blocking output from MBON-α3 impairs pre-exposure memory performance [F(2,28)=10.50, p=0.0004, n=10-11; Tukey’s multiple comparisons test: UAS-*Shi*^ts1^ *vs.* G0239-GAL4, p=0.7412; UAS-*Shi*^ts1^ *vs.* G0239-GAL4;UAS-*Shi*^ts1^, p=0.0043; G0239-GAL4 *vs.* G0239-GAL4;UAS-*Shi*^ts1^, p=0.0006]. (**C**) Odor pre-exposure reduces trained odor responses of MBON-γ2α′1 compared to a novel odor [for MCH: W(7)=24, p=0.0436; for OCT: t(7)=2.689, p=0.0311] (**D**) Blocking output from MBON-γ2α′1 impairs pre-exposure memory performance [F(2,45)=5.063, p=0.0104, n=16; Tukey’s multiple comparisons test: UAS-*Shi*^ts1^ *vs.* MB077B-GAL4, p=0.9823; UAS-*Shi*^ts1^ *vs.* MB077B-GAL4;UAS-*Shi*^ts1^, p=0.0284; MB077B-GAL4 *vs.* MB077B-GAL4;UAS-*Shi*^ts1^, p=0.0181]. (**E**) Restricting MBON block to the test phase using GtACR1 reveals a requirement for MBON-γ2α′1 but not MBON-α3 to express the pre-exposure memory [F(4,50)=7.188, p=0.0001, n=11; Šídák’s multiple comparisons test: UAS-GtACR1 *vs.* G0239-GAL4, p=0.9435; UAS-GtACR1 *vs.* G0239-GAL4;UAS-GtACR1, p=0.0337; G0239-GAL4 *vs.* G0239-GAL4;UAS-GtACR1, p=0.2668; UAS-GtACR1 *vs.* MB077B-GAL4, p=0.9249; UAS-GtACR1 *vs.* MB077B-GAL4;UAS-GtACR1, p=0.0001; MB077B-GAL4 *vs.* MB077B-GAL4;UAS-GtACR1, p=0.0027]. (**F**) Blocking MBON-α3 during memory testing impairs the expression of LI of appetitive memory. The inhibitory effect of odor X pre-exposure on appetitive memory is not observed when α3 MBONs are blocked at retrieval [F(2,27)=5.063, p=0.0136, n=10; Tukey’s multiple comparisons test: UAS-GtACR1 *vs.* G0239-GAL4, p=0.9965; UAS-GtACR1 *vs.* G0239-GAL4;UAS-GtACR1, p=0.0248; G0239-GAL4 *vs.* G0239-GAL4;UAS-GtACR1, p=0.0297]. All behavioural pre-exposure experiments were performed in the context A (clear tubes) for pre-exposure and testing phases.

### α3 MBON output is required for expression of latent inhibition of appetitive memory

We last tested whether the activity of the γ2α′1 and α3 MBONs was required for the expression of LI. Flies expressing UAS-*GtACR1* in γ2α′1 and α3 MBONs were subjected to the standard pre-exposure regimen followed by appetitive conditioning (Fig. 5F). Blocking output from γ2α′1 MBONs impaired the expression of appetitive memory, so their contribution to LI could not be further tested (data not shown). However, silencing the α3 MBONs during testing by illuminating the flies with green light, abolished the effect of LI that was observed in the control flies.

## Discussion

In this study we demonstrate latent inhibition (LI) in *Drosophila* and identify an underlying neuronal mechanism. We find that repeated odor presentation forms a labile self-reinforced aversive memory for that odor, which can temporarily compete with the expression of a newly acquired appetitive memory for that same odor. During memory testing the conditioned odor should therefore activate both the memory of the pre-exposure (odor-self) and that of the appetitive conditioning (odor-sugar). Importantly, the aversive pre-exposure memory is labile which means the LI effect is transient. As a result, the appetitive memory performance exhibits ‘spontaneous recovery’. These results demonstrate that a retrieval (R) model underlies LI in the fly. An acquisition (A) model is not supported because flies acquire an associative reward memory for the odor after pre-exposures of that odor. Instead, the expression of the learned approach performance is impeded by the co-expression of a competing aversive pre-exposure memory. In further support of this R model, the same pre-exposure regimen caused facilitation of a subsequently acquired aversive olfactory memory.

In this instance the pre-exposure memory adds to the new aversive associative memory, rather than competes with an appetitive memory.

A defining feature of LI is a sensitivity to the consistency of the context in which the pre-exposure, learning and testing are carried out ^61^. Changing between the clear and paper-lined tubes did not impair LI, suggesting that the flies likely consider these to be a similar context. However, if odor pre-exposure, learning and testing were performed in different contexts (ie. a copper grid-lined versus a paper-lined, or clear tube) LI was abolished. Most strikingly, LI could be restored if copper grid tubes were used to provide the same context during pre-exposure and testing. In line with prior theories and studies of LI in other animals ^62,63^, these results suggest that flies learn an association between the odor and the context in which it is experienced during the non-reinforced pre-exposure. As a result, the pre-exposure memory gains context-dependence, and our experiments show it is not retrieved if the context is different when memory is tested. The failure to retrieve the pre-exposure memory in a different context manifests as a loss of LI - the appetitive memory is fully expressed. Our study therefore reveals that the context-dependency of LI results from the ability (correct context, LI evident), or inability (wrong context, no LI), to retrieve the pre-exposure memory. In addition, context only plays a role in the expression of the pre-exposure memory when it is in conflict with a subsequently acquired appetitive memory. Further work will be required to define what the flies recognise as a ‘change of context’. There are many possibilities including, background odors, tube/paper texture, relative luminance, and other flies in the group.

We found that the odor-driven activity of γ2α′1 and α3 DANs increased with repeated odor pre-exposure and that they were required for the formation of the odor pre-exposure memory. In addition, the odor-specific responses of the corresponding MBONs were depressed following pre-exposure. We therefore conclude that ramping odor-driven DAN activity assigns negative value to the odor itself by depressing odor-specific KC connections onto the γ2α′1 and α3 MBONs. In support of this model, repeated pre-exposure of flies to a lower and less innately aversive odor concentration did not increase the activity of the α3 DANs, or form an aversive pre-exposure memory. Importantly, reduced odor-activation of the approach-directing γ2α′1 and α3 MBONs MBONs is sufficient to account for the aversive nature of pre-exposure memory. Moreover, both expression of pre-exposure memory and LI are abolished if the α3 MBONs are blocked during testing, confirming the model that LI is produced by the expression of the aversive pre-exposure memory competing with that of the associative reward memory.

Prior work has described odor-driven activity of the PPL1-α′3 DANs, and subsequent depression of odor-specific responses of the α′3 MBONs to underlie how flies become familiar with an odor following repeated short exposures ^17^. In contrast, we show that two longer and spaced odor exposures produce an aversive memory that manifests as plasticity of γ2α′1 and α3 DANs and MBONs. Moreover, whereas we show a retrieval defect underlies LI, a reduced attention/familiarity to the odor following pre-exposures would be expected to result in a subsequent acquisition defect.

LI has often been compared to memory extinction ^64^ and our work in the fly shows that very similar neuronal mechanisms account for both of these phenomena. Pre-exposure learning in *Drosophila* appears to follow similar rules to extinction learning following aversive olfactory conditioning; 2 spaced trials with 15 min ITI is more efficient than massed training with 1 min ITI ^33^ and in both cases a resulting parallel opposing odor-nothing memory inhibits the retrieval/expression of the odor-punishment or odor-reward memory ^33,65^. The obvious difference is that the interfering non-reinforced odor memory is formed before learning for LI, and after learning for extinction.

Our studies of learning, extinction and LI suggest that flies acquire and store all of their experience (rewarded and unrewarded, punished and unpunished) as parallel memory traces ^33–35,65,66^. As a result, when evoked by an appropriate cue, the relevant experiences are compared/combined at the time of retrieval to determine the most fitting behavioral outcome. Such a model is reminiscent of the Miller and Matzel ^67^ comparator hypothesis, devised mostly from experiments in rodents ^11^. Since recent studies suggest similar processes underlie extinction of fear in flies, rodents and humans ^33,68–72^, it seems likely that the mechanism of LI we describe here will also be relevant across phyla.

## Material and Methods

### Fly stock maintenance and fly strains

All *Drosophila melanogaster* strains were reared at 25 °C and 40-50% humidity, except where noted, on standard cornmeal-agar food in 12:12 h light:dark cycle. Canton-S flies were used as wild-type (WT). Transgenes were expressed with previously described GAL4 lines: MB247 ^43^, R58E02-GAL4 ^29^, MB504B-GAL4, MB296B-GAL4, MB058B-GAL4, MB630B-GAL4 and MB077B-GAL4 ^73,74^, c061-GAL4;MBGAL80 ^75^ and G0239-GAL4 ^76^. For behavioral experiments UAS-*Shi*^ts1^ ^48^, UAS-CsChrimson::mVenus ^77^ and UAS-GtACR1 ^49^ were expressed under the control of the respective GAL4–line. For the live-imaging experiments UAS-GCaMP6m ^78^ was expressed with the respective GAL4. We used mixed sex flies for behavior and imaging experiments.

### Behavioral experiments

Male flies from the GAL4 lines were crossed to UAS-*Shi*^ts1^, UAS-CsChrimson or UAS-GtACR1 females, except in the case of the c061-GAL4;MBGAL80 crosses where UAS-*Shi*^ts1^ males were crossed to c061-GAL4;MBGAL80 females. For heterozygous controls GAL4 or UAS-*Shi*^ts1^ flies were crossed to WT. All flies were raised at 25 °C and mixed sex populations of 4–8-day-old flies were used in all experiments. Approximately 80-100 flies were placed in a 25 ml vial containing 1% agar (as a water source) and a 20 × 60 mm piece of filter paper for 18–24 h before training and were kept starved for the entire experiment. For experiments to evaluate the requirement for hunger, flies were fed *ad libitum* by housing them in a 25 ml vial containing standard food for 14–22 h before behavioral experiments. Prior to optogenetic experiments all flies were housed on standard cornmeal food supplemented with 1 mM retinal for 3 days, before being starved as described above. For experiments involving neuronal blockade with UAS-*Shi^ts1^*, the time courses of the temperature shifting are provided alongside each graph of memory performance. For *Shi^ts1^* experiments, flies were transferred to the restrictive 32°C 30 min before the targeted time, except where noted, to allow for acclimatization to the new temperature. All behavioral experiments were performed using a standard T-Maze and 4-methylcyclohexanol (MCH) and 3-octanol (OCT) diluted in mineral oil (MO) were used as the odors.

For repeated odor exposure/odor pre-exposure experiments an odor X (1:10^−6^ or 1:10^−3^ diluted in MO) or mineral oil (MO) was presented twice for 2 min with an inter trial interval (ITI) of 15 min.

When assessing LI, odor pre-exposure was followed by appetitive olfactory conditioning, performed essentially according to Krashes and Waddell ^39^: flies were exposed for 2 min to odor Y without reinforcement, in a tube with dry filter paper (the conditioned stimulus -, CS-), 30 s of clean air, then 2 min with odor X with saturated 5.8M sucrose, dried on a filter paper (the conditioned stimulus+, CS+). For assessing facilation of aversive memory, odor pre-exposure was followed by aversive olfactory conditioning ^59,79^: flies recieved 1 min with odor X paired with twelve 90 V electric shocks at 5 s intervals (CS+), 45 s of clean air, 1 min with odor Y without reinforcement.

When experiments involved neuronal inhibition using GtACR1, the presentation of odors during the pre-exposure or testing phase was paired with continuous green light (three high-power LEDs [700 mA, centered at 530 nm] were mounted on one arm of the T-maze). Immediately after the second exposure, except where noted, flies are given the choice (2 min in darkness) between the pre-exposed odor A and an alternative odor B (novel odor). Performance Index was calculated as the number of flies in the odor A arm minus the number in the odor B arm, divided by the total number of flies ^59,79^.

When experiments involved neuronal activation using CsChrimson, the presentation of the CS+ odor during the training phase was paired with the presentation of red light (three high-power LEDs [700 mA, centered at 630 nm] were mounted on one arm of the T-maze and triggered for 1 ms at 500 Hz), and the electric shocks were omitted.

Memory performance was assessed by testing flies for their odor-preference between the CS- and the CS+ odors for 2 min in darkness (except where noted). Performance Index was calculated as the number of flies in the CS+ arm minus the number in the CS− arm, divided by the total number of flies. For all behavioral experiments, a single sample, or n, represents the average performance score from two reciprocally trained groups of flies.

Naïve avoidance was performed as essentially as described ^58^. An important alteration of the protocol was to use T-maze tubes lined with filter paper and to replace the filter papers and clean the tubes in between experiments, so that odor concentrations did not accumulate in the tubes. This was particularly important to observe approach towards lower 10^−6^ concentrations. Untrained flies were given 2 min in darkness to choose between a diluted odor (OCT or MCH, either 1:10^−6^ or 1:10^−3^) and air bubbled through mineral oil in the T-Maze. Performance Index was calculated as the number of flies in the odor arm minus the number in the air arm, divided by the total number of flies.

### Two-Photon Calcium Imaging

All flies were raised at 25 °C and 3–8 day-old male and female flies were used in all experiments. Imaging experiments were performed essentially as described previously ^33,34,58,59^. In brief, flies were immobilized on ice and mounted in a custom-made chamber allowing free movement of the antennae and legs. The head capsule was opened under room temperature carbogenated (95% O2, 5% CO2) buffer solution (103 mM NaCl, 3 mM KCl, 5mM N-Tris, 10 mM trehalose, 10 mM glucose, 7mM sucrose, 26 mM NaHCO3, 1mM NaH2PO4, 1.5 mM CaCl2, 4mM MgCl2, osmolarity 275 mOsm, pH 7.3) and the fly, in the recording chamber, was placed under the Two-Photon microscope (Scientifica).

Flies were exposed to odors under the microscope using essentially the same regimens and odor concentrations as those in the behavioral experiments. Flies were subjected to a constant air stream, carrying vapor from mineral oil solvent (air). For the odor pre-exposures, an odor stream was added to the air for 2 min. Flies in the custom chamber were then removed from the microscope and rested for 15 min until being returned to the microscope and given the 2^nd^ odor exposure. The carbogenated buffer was changed before each re-exposure. To emulate the testing phase, after the 2^nd^ exposure, the flies were sequentially exposed to the re-exposed odor and a novel odor, each for 5 s, interspersed by 30 sec of air. As in the behavior experiments the odors were MCH and OCT, and they were used reciprocally. GCaMP responses were measured in the relevant DANs and MBONs during pre-exposure and test phases.

One hemisphere of the brain was randomly selected to image the dendritic field of each MBON and the presynaptic terminals of each DAN. Flies that did not respond to one of the two presented odors were excluded from the analyses in this study. Each n corresponds to a recording from a single fly.

Fluorescence was excited using ~140 fs pulses, 80 MHz repetition rate, centered on 910 nm generated by a Ti-Sapphire laser (Chameleon Ultra II, Coherent). Images of 256 x 256 pixels were acquired at 5.92 Hz, controlled by ScanImage 3.8 software ^80^. Odors were delivered using a custom-designed system ^81^.

For analysis, two-photon fluorescence images were manually segmented using Fiji ^82^. Movement of the animals was small enough such that images did not require registration. For subsequent quantitative analyses, custom Fiji and MATLAB scripts were used. The baseline fluorescence, F_0_, was defined for each stimulus response as the mean fluorescence F from 2 s before and up to the point of odor presentation. F/F_0_ accordingly describes the fluorescence relative to this baseline. For the imaging data, the area under the curve (AUC) was measured as the integral of F/F_0_ during the 5 s odor stimulation.

### Statistical Analysis

Statistical analyses were performed in GraphPad Prism. All behavioral data were analyzed with an unpaired t-test or a one-way ANOVA followed by a posthoc Tukey’s or Bonferroni’s multiple comparisons test. No statistical methods were used to predetermine sample size. For the imaging experiments odor-evoked responses were compared by a paired t-test for normally distributed data, otherwise a Wilcoxon matched-pairs signed rank test was used for non-Gaussian distributed data. Normality was tested using the Shapiro-Wilk normality test. For imaging data, a method for outlier identification was run for each dataset (ROUT method), which is based on the False Discovery Rate (FDR). The FDR was set to the highest Q value possible (10%). In datasets in which potential outliers were identified, statistical analyses were performed by removing all odor-evoked responses for those flies. The analyses with or without the outliers were not different, so we decided to maintain and present the complete datasets, which may contain potential outliers.

## Author contributions

Designed research P.F.J., Z.O., P.V-G., J.F., S.W., Performed research P.F.J., Z.O., P.V-G., J.F., S.V-M., Analyzed data P.F.J, Z.O., J.F., P.V-G., S.V-M. Resources S.W. Writing S.W., P.F.J., Z.O., J.F. Supervision S.W. Funding Acquisition S.W.

## Acknowledgments

We thank G. Rubin, FlyLight, B. Dickson and the Bloomington Stock Center for flies. We are grateful to Geraldine Wright and members of the Waddell group for discussion. P.V-G. is funded by the Sloane Robinson/Clarendon Scholarship given by the University’s Clarendon Fund scheme and Keble College and from the Consejo Nacional de Ciencia y Tecnología (CONACYT). S.W. is funded by a Wellcome Principal Research Fellowship (200846/Z/16/Z) and an ERC Advanced Grant (789274).

## Notes

Conflict of Interest: The authors declare no competing financial interests.

### Competing Interest Statement

The authors have declared no competing interest.

